# Bayesian spatial prediction of three medically important tick species in Illinois

**DOI:** 10.64898/2026.04.16.719082

**Authors:** Abrar Hussain, Lelys Bravo de Guenni, Nohra Mateus-Pinilla, Rebecca L. Smith

**Author notes:** **Corresponding Author** Abrar Hussain, Room 2533, 2001 S Lincoln Ave, Urbana, IL, USA 61802.

## Abstract

Tick-borne diseases are reported across much of Illinois, yet tick surveillance remains spatially uneven, leaving many counties with limited species-specific observations. We developed Bayesian hierarchical spatial models to predict county-level relative abundance of *Amblyomma americanum, Dermacentor variabilis*, and *Ixodes scapularis* using surveillance data from 2019-2022. Relative abundance was defined as the species-specific share of the collected focal ticks and modeled using beta-binomial models that incorporated climate, land cover, forest fragmentation, deer habitat suitability, and BYM2 spatial effects in INLA. Spatial five-fold cross-validation evaluated whether environmental information improved prediction beyond spatial structure alone. *A. americanum* showed the highest predicted relative abundance in southern Illinois, with additional elevated areas in central-western counties, and deer habitat suitability was its clearest environmental predictor. *D. variabilis* was broadly distributed and best represented by the spatial-only model. *I. scapularis* was concentrated mainly in northern Illinois, with the climate providing the strongest environmental signal. Statewide predictions extended inference to counties lacking species-specific observations and identified where uncertainty remained greatest, providing a basis for prioritizing additional surveillance across Illinois.

## Introduction

Medically important ixodid ticks are expanding their geographic ranges across much of the United States, reshaping the landscape of vector-borne disease risk (Alkishe et al. 2021). In Illinois, three species are of particular concern for human and animal health: the lone star tick (*Amblyomma americanum*), the American dog tick (*Dermacentor variabilis*), and the blacklegged tick (*Ixodes scapularis*) (Carson et al. 2022). These species transmit pathogens associated with Lyme disease, anaplasmosis, ehrlichiosis, spotted fever group rickettsioses, and babesiosis, while bites from some species, particularly *A. americanum*, are also associated with emerging conditions such as alpha-gal syndrome (Jongejan and Uilenberg 2004; Lankester et al. 2007; Goddard and Varela-Stokes 2009; Diuk-Wasser et al. 2010; Brown et al. 2011; Scott Dahlgren et al. 2016; Eisen et al. 2016; Soucy et al. 2018). Recent surveillance and case data indicate that tick-borne diseases (TBDs) have been reported from almost every county in Illinois and that the incidence of several TBDs has increased in recent years (Chakraborty et al. 2025), underscoring the need for spatially explicit estimates of vector distributions at sub-state scales.

Tick surveillance relies on a combination of active sampling (e.g., drag/flagging, CO_2_ traps) and passive submissions from the public, veterinarians, and health agencies (Eisen and Paddock 2021; Chakraborty et al. 2025). Because these activities are logistically demanding and resource-intensive, sampling intensity is often uneven across space and time (Guillot et al. 2023), leaving many counties with sparse or no tick surveillance data. A previous statewide analysis identified substantial geographic gaps in available tick records, with many Illinois counties having limited or no surveillance data (Hussain et al. 2025a). Such sparsity limits inference about relative abundance in poorly sampled counties and makes it difficult to identify where additional surveillance is most needed. Spatial models can help address these limitations by borrowing information across locations, generating predictions for counties without species-specific observations, and quantifying uncertainty in predicted relative abundance (Sharma et al. 2024).

The distribution and abundance of *A. americanum, D. variabilis*, and *I. scapularis* are influenced by climate, habitat, and host availability (Bacon et al. 2022). Temperature and precipitation patterns shape tick survival, development, and activity (Burtis et al. 2016), while land cover, especially forest, grassland, and ecotonal habitats, affects microclimate and encounter rates with vertebrate hosts (González et al. 2023; Di et al. 2023). White-tailed deer can contribute to the maintenance and dispersal of several medically important tick species and may influence their distributions across heterogeneous Midwestern landscapes (Huang et al. 2019; Tardy et al. 2023; Rochlin et al. 2025). At finer scales, forest fragmentation and composition can create a patchwork of high- and low-risk areas that are not captured by climate alone (Brownstein et al. 2005). However, the predictive importance of these environmental and host-related factors may differ among tick species, underscoring the need to determine whether they improve prediction beyond residual spatial structure alone. Bayesian hierarchical spatial models are well-suited to this problem because they can account for residual spatial dependence, extend inference into data-poor areas, and quantify uncertainty in spatial predictions (Michal et al. 2025). Integrated Nested Laplace Approximation (INLA) provides computationally efficient inference for such areal models (Bakka et al. 2018).

In this study, we developed Bayesian spatial models of county-level relative abundance for *A. americanum, D. variabilis*, and *I. scapularis* in Illinois. Our objectives were to predict relative abundance across all counties, including those lacking species-specific observations; quantify geographic variation in prediction uncertainty; and determine whether climate, land cover, forest fragmentation, and deer habitat suitability improve prediction beyond residual spatial structure alone. This framework was designed to identify surveillance gaps and areas where additional sampling could most improve understanding of tick distributions.

## Materials and Methods

### Study area and spatial units

The study was conducted across Illinois, which spans substantial geographic gradients in climate, land cover, habitat structure, and vertebrate host communities (Illinois Department of Natural Resources 2025). Counties were used as the areal units for all analyses; the shapefile for Illinois counties was obtained from the Illinois Natural Resources Geospatial Data Clearinghouse (Illinois State Geological Survey 2025). Spatial data were processed in R Statistical Software (version 4.5.2) (R Core Team 2025), and all spatial operations were performed using simple features (**sf**) (Pebesma 2025) and raster-based packages (**terra, raster**) (Hijmans 2025b; Hijmans 2025a), with projections chosen to balance interpretability (geographic coordinates, EPSG:4326) and metric calculations (projected coordinates, UTM Zone 16N, EPSG:26916) (EPSG.io 2025).

### Surveillance data preparation

County-level surveillance data for *A. americanum, D. variabilis*, and *I. scapularis* were compiled for 2019-2022 using the same statewide surveillance dataset and data-curation criteria as previous analysis of medically important ticks in Illinois (Hussain et al. 2025a). Because the surveillance records did not consistently distinguish true species non-detection from the absence of sampling information, counties without reliable species-specific observations were retained as missing (NA) rather than interpreted as confirmed absences, consistent with the corrected treatment in the previous analysis (Hussain et al. 2025b).

To account for variation in the number of ticks represented in the surveillance records, we expressed the response as species-specific relative abundance. For species ***s*** in county ***i***, relative abundance was defined as the number of ticks of that species divided by the total number of the three focal tick species represented in that county, yielding a proportion between 0 and 1. In the Bayesian models, the original integer species count was retained as the response, and the total number of collected ticks was specified as the beta-binomial denominator ***N***_*i*_ Thus, the modeled probability represents the expected relative abundance of a species among collected ticks, rather than absolute tick density or abundance per unit sampling effort, with total ticks serving as an indirect measure of sampling effort (Figure 1).

**Figure 1.**
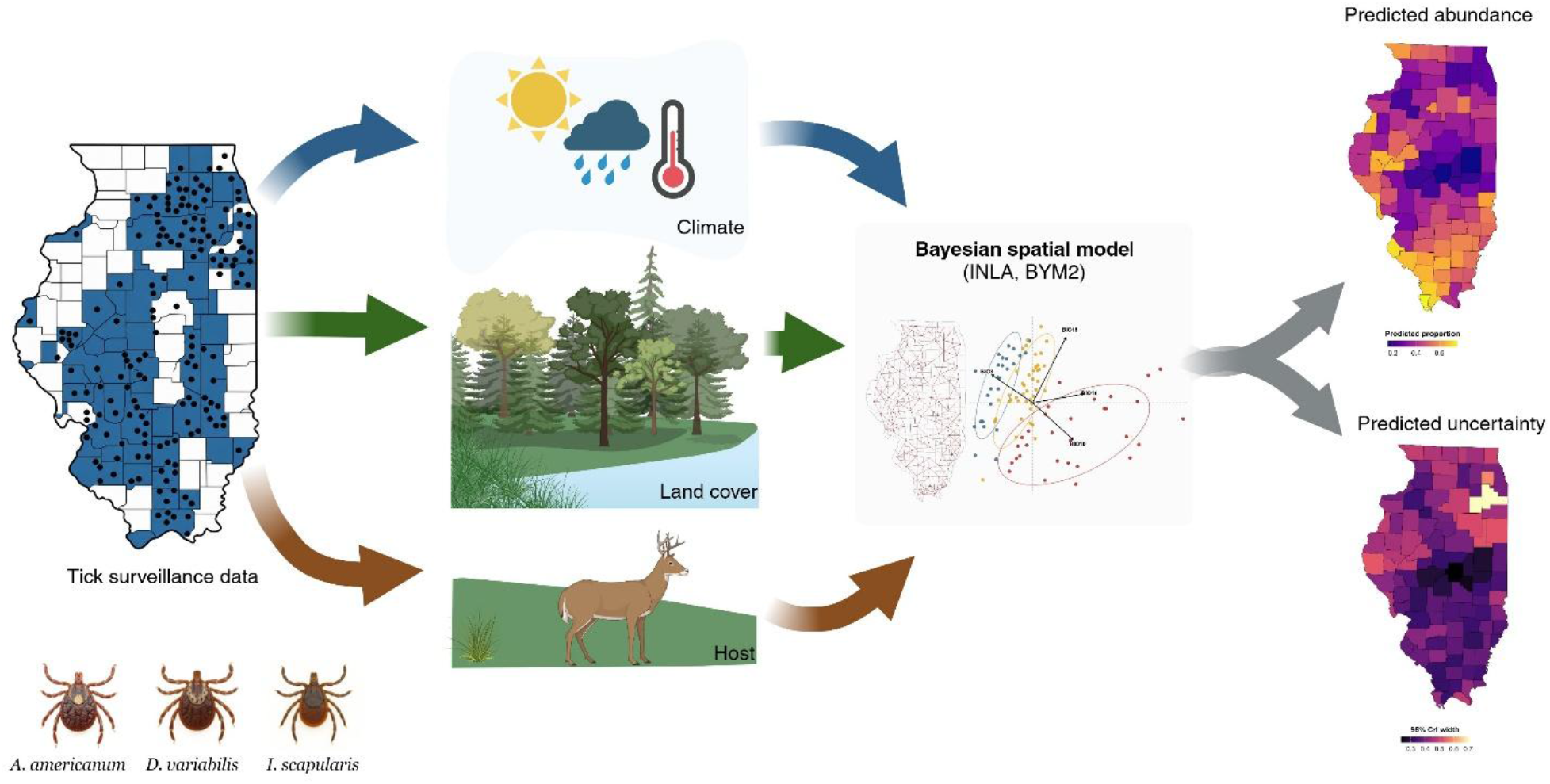
Conceptual framework of the Bayesian spatial modeling approach used to predict county-level tick relative abundance and quantify prediction uncertainty across Illinois.

### Environmental covariates

Climate data were obtained from the WorldClim 2.1 historical CRU-TS 4.09 dataset at approximately 2.5-arc-minute resolution (WorldClim 2025). Monthly minimum temperature, maximum temperature, and precipitation layers for 2019-2022 were used to derive annual bioclimatic variables. Four biologically relevant variables were retained: mean temperature of the wettest quarter (BIO8), mean temperature of the warmest quarter (BIO10), precipitation of the wettest quarter (BIO16), and precipitation of the warmest quarter (BIO18). County-level means were extracted for each year and averaged across 2019-2022 to produce a single climate profile per county. Land-cover characteristics were derived from the 2019 National Land Cover Database (NLCD) (U.S. Geological Survey 2025). County-level forest, grassland, and wetland cover were summarized together with a forest-fragmentation index derived using the **landscapemetrics** package (Hesselbarth et al. 2025). The fragmentation index combined standardized patch density, mean patch area, edge density, and clumpiness, with higher values representing greater fragmentation. To incorporate host-related habitat suitability, we used white-tailed deer habitat quality scores from the Deer Land Cover Utility (LCU) dataset developed by (Mori et al. 2024) and published in the Illinois Data Bank (Illinois Data Bank 2025).

Pairwise correlations among the selected environmental covariates were examined to characterize correlation structure among the climate and land-cover variables (Figure S1). Because several variables were correlated, principal component analysis (PCA) was applied separately to the climate and land-cover datasets to reduce dimensionality and collinearity before model fitting.

### Principal component analysis

PCA was applied separately to the four climate variables and four land-cover variables using the *prcomp()* function in **R** after centering and scaling. Climate PC1 and PC2 were retained and together explained 82.49% of climate variation. Land-cover PC1-PC3 were retained and together explained 86.42% of land-cover variation. Deer habitat suitability remained a separate predictor. Retained PCA scores and deer suitability were standardized to a mean of 0 and a standard deviation of 1 before model fitting.

### Bayesian spatial modeling, implementation, and model selection

Species-specific relative abundance was modeled using Bayesian beta-binomial regression implemented with INLA (Bakka et al. 2018). For species ***s*** in county ***i***, the observed species count was modeled conditional on the total number of focal ticks represented in that county. A logit link related the expected species-specific relative abundance to environmental covariates and, where applicable, a spatial random effect. The beta-binomial likelihood allowed for extra-binomial variation in species composition.

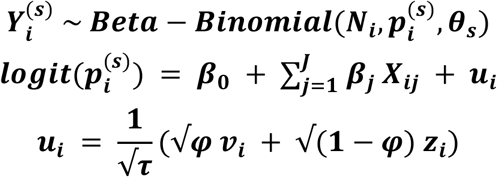

Here, ***Y*** is the observed species count, ***N*** is the total number of focal ticks represented in the county, ***p*** is the expected relative abundance, ***θ*** is the species-specific beta-binomial overdispersion parameter, ***X*** denotes the standardized environmental predictors, and u is the BYM2 spatial effect. In the BYM2 parameterization, ***τ*** is the overall precision, ***v*** is the scaled spatially structured component, ***z*** is the unstructured component, and ***φ*** controls the relative contribution of structured spatial variation.

A symmetric five-nearest-neighbor (KNN-5) graph was constructed from representative county points in UTM Zone 16N and used for the BYM2 component. Six candidate models were fitted separately for each species: M1, full environmental effects without BYM2; M2, spatial-only BYM2; M3a, climate PCs + BYM2; M3b, land-cover PCs + BYM2; M3c, deer habitat suitability + BYM2; and M3d, full environmental effects + BYM2. The full environmental set included climate PC1-PC2, land-cover PC1-PC3, and deer habitat suitability.

Candidate models were compared using the Watanabe-Akaike information criterion (WAIC), deviance information criterion (DIC), and log conditional predictive ordinate (LCPO), with lower values indicating better model support. A final model was selected separately for each species, and PIT and CPO diagnostics were examined for the selected models. Posterior fixed effects were summarized as odds ratios with 95% credible intervals, and associations were considered credibly different from zero when the corresponding 95% credible interval on the regression-coefficient scale excluded zero.

### Spatial cross-validation

Spatial predictive performance of the best-fit model for each tick species was evaluated using balanced five-fold cross-validation constructed separately for each species from counties with available species-specific observations. Initial geographic clusters were generated from standardized projected county coordinates using k-means clustering, and counties were reassigned as needed to ensure fold sizes differed by no more than 1 county while preserving geographic coherence. For each fold, outcomes in the held-out counties were set to missing, the model was refitted to the remaining observations, and posterior mean relative-abundance estimates were extracted for the held-out counties. Each observed county, therefore, contributed one out-of-fold prediction. Predictive performance was summarized using mean squared error (MSE), root mean squared error (RMSE), mean absolute error (MAE), and mean prediction bias. Predictive uncertainty was evaluated using 95% posterior predictive intervals generated from the beta-binomial posterior predictive distribution for each held-out county, based on its original total focal-tick count. Empirical coverage was calculated as the proportion of observed relative-abundance values falling within these intervals.

To assess whether environmental covariates improved prediction beyond spatial structure alone, identical folds were used to compare the best-fit model with a spatial-only model and, if the spatial-only model was the best fit, with an environmental-plus-spatial model. Relative changes in RMSE and MAE were used to quantify improvement or deterioration in geographically out-of-sample prediction.

### Spatial prediction and uncertainty

The selected model for each species was refitted using all available species-specific observations. Counties without species-specific observations remained missing outcomes but retained environmental covariates and membership in the spatial graph, allowing posterior estimates to be generated for all Illinois counties. Posterior marginals were transformed from the logit scale to the 0-1 relative-abundance scale. For each county, posterior means and 95% credible intervals were extracted, and the width of the 95% credible interval was used to characterize geographic variation in posterior uncertainty.

## Results

### Descriptive surveillance patterns

Species-specific relative-abundance observations were available for 49 counties for *A. americanum*, 76 for *D. variabilis*, and 30 for *I. scapularis*, leaving 53, 26, and 72 counties, respectively, without species-specific observations. Among counties with available observations, median relative abundance was 0.52 for *A. americanum*, 0.64 for *D. variabilis*, and 0.23 for *I. scapularis*. The distributions of observed county-level relative abundance differed among species (Figure S2). Observed relative abundance showed distinct spatial patterns, with *A. americanum* concentrated primarily in southern and central Illinois, *D. variabilis* more broadly distributed across the state, and *I. scapularis* observed in fewer, more spatially heterogeneous counties (Figure 2).

**Figure 2.**
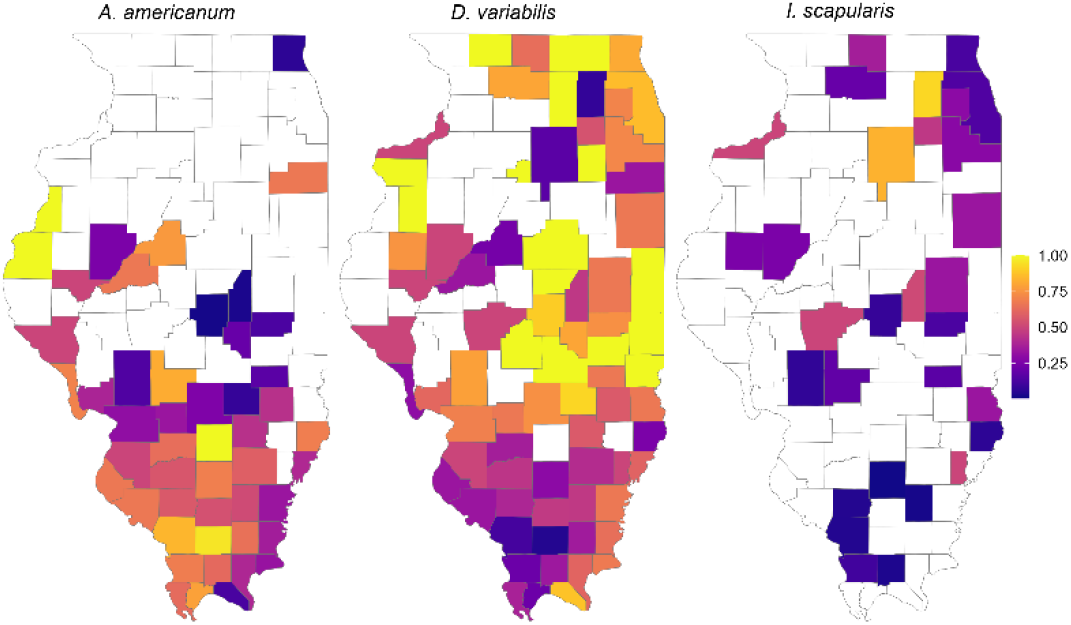
Observed relative abundance of *Amblyomma americanum, Dermacentor variabilis*, and *Ixodes scapularis* across Illinois counties.

### Environmental covariates and principal components

The environmental and ecological covariates showed pronounced spatial variation across Illinois, with distinct climatic, land-cover, fragmentation, and deer-suitability gradients across northern, central, and southern counties (Figure 3). Several covariates were correlated (Figure S1), and PCA summarized these correlated environmental gradients.

**Figure 3.**
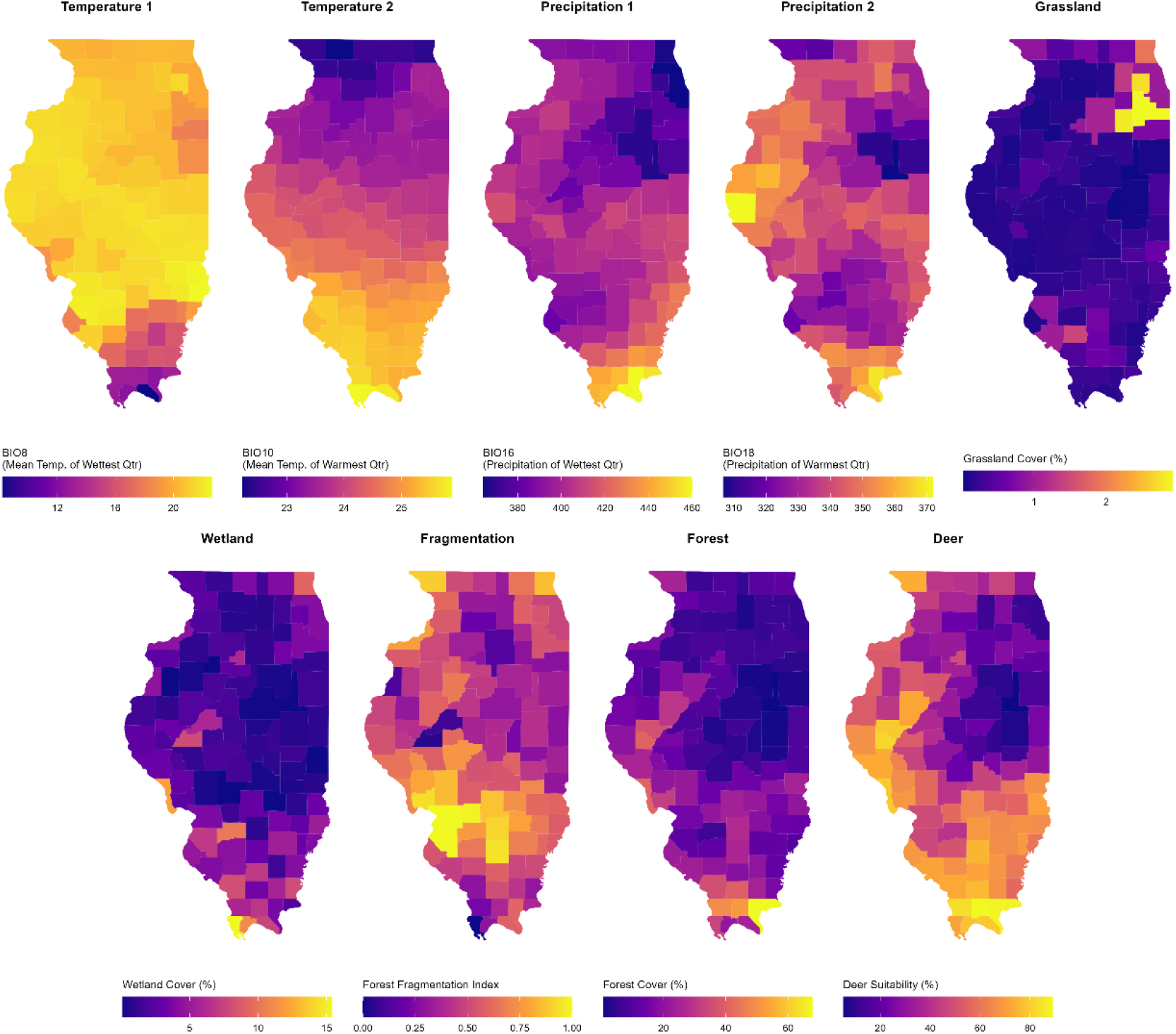
Spatial distribution of the nine environmental and ecological covariates used in the analysis across Illinois counties: BIO8, BIO10, BIO16, BIO18, forest cover, grassland cover, wetland cover, forest fragmentation, and deer habitat suitability.

Climate PC1 explained 60.20% of the variation and represented the main north-south climatic gradient, with higher scores associated with warmer warmest-quarter temperatures (BIO10) and greater wettest- and warmest-quarter precipitation (BIO16 and BIO18), and lower wettest-quarter temperatures (BIO8). Scores were generally higher in southern Illinois, lower in the north, and intermediate in the central region. Climate PC2 explained 22.29% of the variation and primarily represented warm-season precipitation and temperature gradients (Figure 4; Table S2).

**Figure 4.**
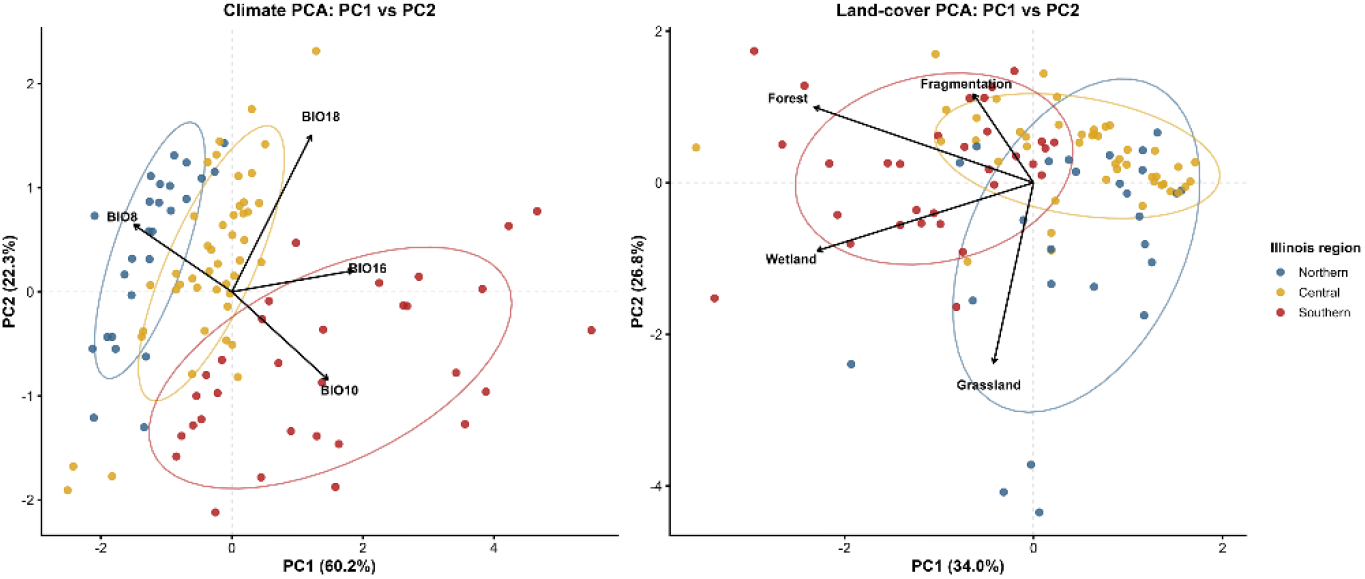
Principal component scores for the first two climate and land-cover components across Illinois counties. Climate PC1-PC2 summarizes the dominant climatic gradients, while land-cover PC1-PC2 summarizes the major ecological gradients.

For land cover, PC1 explained 34.05% of the variance and was driven predominantly by forest and wetland cover, both loading in the same direction; southern counties generally occupied the forest-wetland end of this gradient. PC2 explained 26.80% and primarily contrasted grassland with fragmentation and, to a lesser extent, forest cover (Figure 4). The first two components explained 60.84% of land-cover variation, increasing to 86.42% with PC3, which was dominated by forest fragmentation and retained in the models (Figure S3; Table S2).

Counties are grouped by northern, central, and southern Illinois to illustrate regional separation along these environmental gradients.

### Bayesian model selection and posterior associations

Model support differed among species (Table S3). For *A. americanum*, the full environmental + BYM2 model (M3d) was best supported (WAIC = 250.92). Deer habitat suitability was positively associated with relative abundance (β = 0.92, 95% CrI: 0.31-1.51; OR = 2.51, 95% CrI: 1.36-4.55), while the remaining climate and land-cover effects were not credibly different from zero. For *D. variabilis*, the spatial-only BYM2 model (M2) had the strongest support (WAIC = 320.14), indicating that the measured environmental covariates did not improve model support. For *I. scapularis*, the climate + BYM2 model (M3a) was selected (WAIC = 137.94), with climate PC1 negatively associated with relative abundance (β = −0.55, 95% CrI: −0.99 to −0.08; OR = 0.58, 95% CrI: 0.37-0.92), whereas climate PC2 was not credibly associated (Figure 5). Posterior mean BYM2 mixing parameters (*ϕ*) were 0.28 for *A. americanum*, 0.68 for *D. variabilis*, and 0.28 for *I. scapularis*; credible intervals were broad, indicating uncertainty in the structured-unstructured variance partition. No CPO failures were observed for the selected models; PIT and −log(CPO) diagnostics are shown in Figure S4.

**Figure 5.**
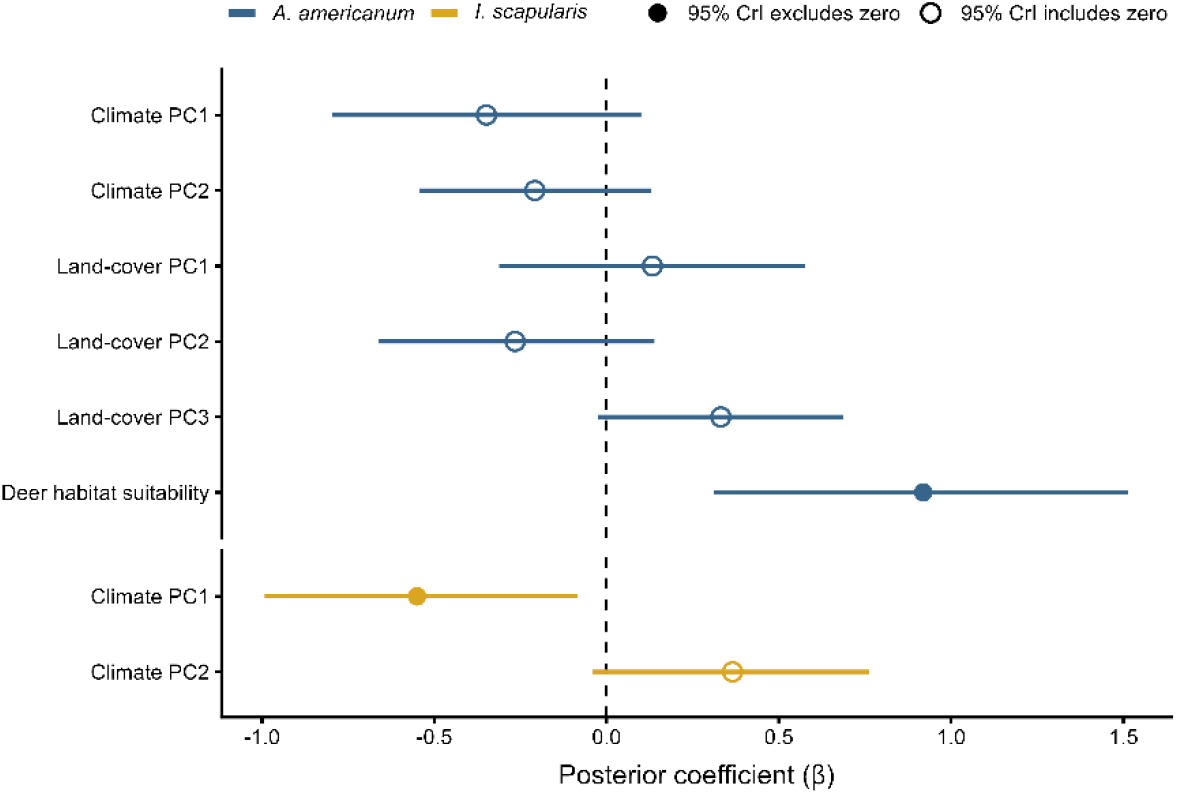
Posterior fixed effect estimates from the selected Bayesian models for *A. americanum* and *I. scapularis*. Points show posterior mean coefficients (β) and horizontal lines show 95% credible intervals; filled points indicate intervals excluding zero. *D. variabilis* is not shown because its selected model was spatial-only.

### Spatial cross-validation

Balanced five-fold spatial cross-validation showed that environmental covariates improved geographically out-of-sample prediction for *A. americanum* and *I. scapularis*, but not for *D. variabilis*. For *A. americanum*, the selected M3d model achieved RMSE = 0.277 and MAE = 0.212, representing reductions of 5.4% and 10.9%, respectively, relative to the spatial-only BYM2 model. For *I. scapularis*, M3a achieved RMSE = 0.236 and MAE = 0.184, improving these metrics by 7.3% and 7.5%. In contrast, the spatial-only model performed best for *D. variabilis* (RMSE = 0.259; MAE = 0.220), whereas adding environmental covariates increased RMSE by 8.3% and MAE by 1.7%. The selected models showed low prediction bias across all three species, with bias values close to zero (Table S4). Empirical coverage of the 95% posterior predictive intervals was 93.9% for *A. americanum*, 97.4% for *D. variabilis*, and 90.0% for *I. scapularis*.

### Spatial prediction and uncertainty

The selected species-specific models generated predicted relative abundance for all 102 Illinois counties, including counties without species-specific observations (Figure 6). Spatial patterns differed among species, with *A. americanum* showing generally higher predicted relative abundance in southern Illinois, *D. variabilis* showing a broader distribution across the state, and *I. scapularis* showing higher predicted relative abundance primarily in northern Illinois. Uncertainty around these predictions was highest in northeastern Illinois for *A. americanum*, west-central Illinois for *D. variabilis*, and east-central Illinois for *I. scapularis*, identifying areas where additional surveillance could most improve prediction precision (Figure S5). Analysis-ready data and R code are available in the GitHub repository listed in the Data Availability statement.

**Figure 6.**
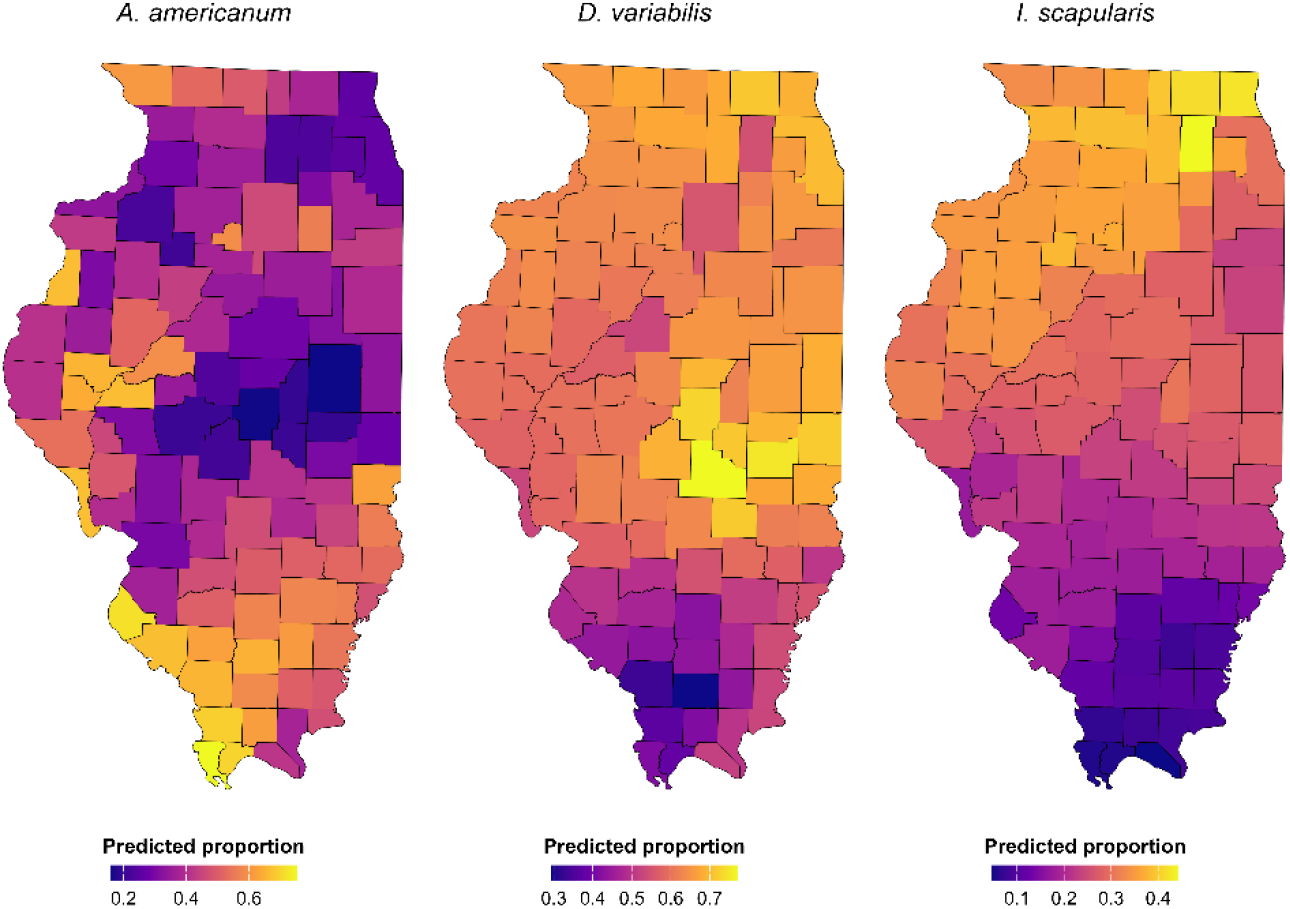
Posterior mean predicted relative abundance of *Amblyomma americanum, Dermacentor variabilis*, and *Ixodes scapularis* across Illinois counties based on the selected species-specific Bayesian models.

## Discussion

This study used Bayesian spatial models to predict county-level relative abundance of three medically important tick species across Illinois, including counties lacking species-specific observations, while evaluating whether environmental and host-related information improved prediction beyond residual spatial structure. Environmental information improved prediction in geographically held-out counties for *A. americanum* and *I. scapularis*, whereas *D. variabilis* was better predicted by the spatial-only model. Deer habitat suitability showed the clearest association with *A. americanum*, while climate emerged as the principal environmental predictor for *I. scapularis*. Together, these models provide statewide predictions across surveillance gaps while identifying where prediction uncertainty is greatest and additional surveillance would be most informative.

### Amblyomma americanum

*A. americanum* showed its highest predicted relative abundance across much of southern Illinois, with additional areas of elevated relative abundance in central-western counties. This pattern is consistent with its historical concentration in the southern Midwest and with reports of a broader distribution farther north and west (Sagurova et al. 2019; Fowler et al. 2022). Range expansion of this species has been linked to warmer conditions, improved overwinter survival, and movement of vertebrate hosts, particularly white-tailed deer (Dawe and Boutin 2016; Rochlin et al. 2022). Similar expansion has been reported in neighboring Midwestern states, including Missouri and Michigan (Louis 2012; Hudman and Sargentini 2018; Fowler et al. 2022). Within Illinois, however, deer habitat suitability emerged as the clearest environmental correlate of relative abundance, suggesting that host-associated habitat conditions may help distinguish areas favorable to *A. americanum* beyond broad climatic and land-cover gradients. This is ecologically plausible because white-tailed deer are important hosts for adult *A. americanum* and can facilitate tick dispersal across landscapes (Dawe and Boutin 2016; Rochlin et al. 2022) . The improvement in spatial cross-validation after incorporating environmental information further indicates that these predictors contributed information beyond residual spatial structure alone. The concentration of predicted relative abundance in southern Illinois also broadly overlaps areas with higher reported incidence of ehrlichiosis and spotted fever group rickettsioses, reinforcing the value of continued surveillance in this region (Chakraborty et al. 2025).

### Dermacentor variabilis

*D. variabilis* remained broadly distributed across Illinois, consistent with its generalist ecology and wide habitat tolerance throughout the eastern United States (Minigan et al. 2018). Similar widespread occurrence has been reported in neighboring states, including Indiana, where the species has been documented in nearly every county, and Wisconsin, where established populations occur across multiple counties (Indiana Department of Health 2025; Wisconsin Ticks 2025). The lack of additional predictive improvement from the measured environmental covariates may therefore reflect the limited ability of broad county-level climatic and landscape gradients to discriminate areas of higher relative abundance for such an ecologically widespread species. *D. variabilis* occupies a range of habitats, including ecotonal and mixed landscapes (Nelder et al. 2022; TickEncounter 2025), while local host availability, vegetation, microclimate, dispersal history, and other unmeasured processes may contribute to geographic variation not captured by the available predictors (Connor et al. 2019; Paradinas et al. 2023).

### Ixodes scapularis

*I. scapularis* showed its highest predicted relative abundance in northern Illinois, with additional elevated areas in selected central counties, consistent with the long-established concentration of this species in northern Illinois and the broader Upper Midwest (Eisen and Eisen 2023a; Dong et al. 2025) . Climate was the clearest environmental signal for this species. Higher relative abundance was associated with the lower end of climate PC1, a gradient characterized by comparatively cooler warmest-quarter temperatures, warmer wettest-quarter temperatures, and lower precipitation during the wettest and warmest quarters. Rather than implicating any single climatic variable, this pattern suggests that the seasonal climate regime characteristic of northern Illinois helps distinguish areas favorable to *I. scapularis*. This interpretation is consistent with evidence that temperature and moisture influence *I. scapularis* development, survival, and seasonal activity across the northern United States (Eisen and Eisen 2023b; Eisen and Eisen 2023a; Price et al. 2024). At the same time, the absence of a clear independent land-cover effect should not be interpreted as evidence that forest habitat is unimportant. Forest cover, leaf-litter conditions, vertebrate hosts, and local moisture strongly influence *I. scapularis* ecology, but these relationships can operate at much finer spatial scales than county-level averages and may therefore be poorly represented by broad landscape gradients (Linske and Williams 2024; Short and Pesapane 2024) . The improvement in geographically held-out prediction when climate was included further indicates that this climatic signal was not simply reproducing the underlying spatial pattern. The northern and central areas of higher predicted relative abundance also broadly coincide with higher reported incidence of Lyme disease and anaplasmosis in Illinois (Chakraborty et al. 2025), highlighting these areas for continued vector and pathogen surveillance.

Across the three species, the contrasting model structures reinforce that medically important ticks in Illinois are not governed by a single set of ecological controls. Previous work in Illinois has likewise demonstrated species-specific climatic associations among these ticks (Bacon et al. 2022). Environmental information improved prediction for *A. americanum* and *I. scapularis*, whereas *D. variabilis* was better predicted by residual spatial structure alone, emphasizing the value of species-specific predictive models. Importantly, the models extended inference to counties lacking species-specific observations, while geographic variation in credible-interval width identified areas where those predictions were more uncertain. Higher uncertainty in northeastern Illinois for *A. americanum*, west-central Illinois for *D. variabilis*, and east-central Illinois for *I. scapularis* highlights areas where additional standardized surveillance could improve prediction precision, particularly given the uneven spatial coverage of tick surveillance in the United States (Mader et al. 2021). Relevant human and canine tick-borne disease cases have also been reported within these broader regions, further increasing the value of clarifying local vector distributions (Chakraborty et al. 2025) .

Several limitations arise from heterogeneous surveillance and county-level aggregation. Collection method, timing, sampling intensity, and detection probability can influence observed tick patterns (Rynkiewicz and Clay 2014; Kessler et al. 2022; Sirén et al. 2024) . Because standardized measures of field-sampling effort were unavailable, species counts were modeled using the total number of focal ticks as the beta-binomial denominator; however, this approach cannot fully account for heterogeneous detection or sampling effort. The BYM2 spatial effect was used to capture residual geographic structure not explained by measured covariates, but county-level aggregation may still obscure finer-scale variation in vegetation, microclimate, and host availability, and can affect apparent entomological-epidemiological relationships.

## Conclusions

By integrating tick surveillance with environmental, host-related, and spatial information, this study generated statewide predictions of relative abundance for three medically important tick species, including counties lacking species-specific observations. The findings revealed distinct species-specific patterns: deer habitat suitability was particularly informative for *A. americanum*, climate improved prediction of *I. scapularis*, whereas *D. variabilis* was better predicted by the spatial-only model. Prediction uncertainty further identified areas where estimates were less precise and additional surveillance would be most informative. Together, these results demonstrate the value of species-specific spatial modeling for extending inference beyond sampled locations and for prioritizing standardized surveillance where additional observations could most improve understanding of tick distributions.

## Acknowledgement

We thank all the researchers who put their efforts into tick collections.

## Credit authorship contribution statement

**Abrar Hussain**: Conceptualization, Methodology, Data Curation, Data visualization, Formal analysis, Software, Writing an original draft, Review & editing; **Nohra Mateus-Pinilla**: Methodology, Review & editing; **Lelys Bravo de Guenni**: Methodology, Formal analysis, Review & editing; **Rebecca L. Smith**: Conceptualization, Review & editing, Supervision.

## Data availability

The relevant codes and data can be accessed here: https://github.com/abrarhussain95/Predicting_IL_tick

## References

Alkishe A, Raghavan RK, Peterson AT (2021) Likely Geographic Distributional Shifts among Medically Important Tick Species and Tick-Associated Diseases under Climate Change in North America: A Review. Insects 2021, Vol 12, Page 225 12:225. 10.3390/INSECTS12030225

Bacon EA, Kopsco H, Gronemeyer P, Mateus-Pinilla N, Smith RL (2022) Effects of Climate on the Variation in Abundance of Three Tick Species in Illinois. J Med Entomol 59:700– 709. 10.1093/JME/TJAB189

Bakka H, Rue H, Fuglstad GA, Riebler A, Bolin D, Illian J, Krainski E, Simpson D, Lindgren F (2018) Spatial modeling with R-INLA: A review. Wiley Interdiscip Rev Comput Stat 10:e1443. 10.1002/WICS.1443

Brown HE, Yates KF, Dietrich G, MacMillan K, Graham CB, Reese SM, Helterbrand WS, Nicholson WL, Blount K, Mead PS, Patrick SL, Eisen RJ (2011) An Acarologic Survey and Amblyomma americanum Distribution Map with Implications for Tularemia Risk in Missouri. Am J Trop Med Hyg 84:411. 10.4269/AJTMH.2011.10-0593

Brownstein JS, Skelly DK, Holford TR, Fish D (2005) Forest fragmentation predicts local scale heterogeneity of Lyme disease risk. Oecologia 2005 146:3 146:469–475. 10.1007/S00442-005-0251-9

Burtis JC, Sullivan P, Levi T, Oggenfuss K, Fahey TJ, Ostfeld RS (2016) The impact of temperature and precipitation on blacklegged tick activity and Lyme disease incidence in endemic and emerging regions. Parasites & Vectors 2016 9:1 9:606-. 10.1186/S13071-016-1894-6

Carson DA, Kopsco H, Gronemeyer P, Mateus-Pinilla N, Smith GS, Sandstrom EN, Smith RL (2022) Knowledge, attitudes, and practices of Illinois medical professionals related to ticks and tick-borne disease. One Health 15:100424. 10.1016/J.ONEHLT.2022.100424

Chakraborty S, Lyons LA, Winata F, Mateus-Pinilla N, Smith RL (2025) Methods of active surveillance for hard ticks and associated tick-borne pathogens of public health importance in the contiguous United States: a comprehensive systematic review. J Med Entomol 62:675–689. 10.1093/JME/TJAF031

Connor T, Viña A, Winkler JA, Hull V, Tang Y, Shortridge A, Yang H, Zhao Z, Wang F, Zhang J, Zhang Z, Zhou C, Bai W, Liu J (2019) Interactive spatial scale effects on species distribution modeling: The case of the giant panda. Scientific Reports 2019 9:1 9:14563-. 10.1038/s41598-019-50953-z

Dawe KL, Boutin S (2016) Climate change is the primary driver of white-tailed deer (Odocoileus virginianus) range expansion at the northern extent of its range; land use is secondary. Ecol Evol 6:6435–6451. 10.1002/ECE3.2316

Di C, Sulkow B, Qiu W, Sun S (2023) Effects of Micro-Scale Environmental Factors on the Quantity of Questing Black-Legged Ticks in Suburban New York. Applied Sciences 2023, Vol 13, Page 11587 13:11587. 10.3390/APP132011587

Diuk-Wasser MA, Vourc’h G, Cislo P, Hoen AG, Melton F, Hamer SA, Rowland M, Cortinas R, Hickling GJ, Tsao JI, Barbour AG, Kitron U, Piesman J, Fish D (2010) Field and climate-based model for predicting the density of host-seeking nymphal Ixodes scapularis, an important vector of tick-borne disease agents in the eastern United States. Global Ecology and Biogeography 19:504–514. 10.1111/J.1466-8238.2010.00526

Dong D, Paskewitz SM, Tsao JI, Schoville SD (2025) Genetic and Landscape Connectivity of Blacklegged Ticks During Range Expansion in Select States of the Midwestern USA. Ecol Evol 15:e72360. 10.1002/ECE3.72360

Eisen L, Eisen RJ (2023a) Changes in the geographic distribution of the blacklegged tick, Ixodes scapularis, in the United States. Ticks Tick Borne Dis 14:102233. 10.1016/J.TTBDIS.2023.102233

Eisen RJ, Eisen L (2023b) Evaluation of the association between climate warming and the spread and proliferation of Ixodes scapularis in northern states in the Eastern United States. Ticks Tick Borne Dis 15:102286. 10.1016/J.TTBDIS.2023.102286

Eisen RJ, Eisen L, Ogden NH, Beard CB (2016) Linkages of Weather and Climate With Ixodes scapularis and Ixodes pacificus (Acari: Ixodidae), Enzootic Transmission of Borrelia burgdorferi, and Lyme Disease in North America. J Med Entomol 53:250–261. 10.1093/JME/TJV199

Eisen RJ, Paddock CD (2021) Tick and Tickborne Pathogen Surveillance as a Public Health Tool in the United States. J Med Entomol 58:1490. 10.1093/JME/TJAA087

EPSG.io (2025) NAD83 / UTM zone 16N - EPSG:26916. https://epsg.io/26916. Accessed 24 Nov 2025

Fowler PD, Nguyentran S, Quatroche L, Porter ML, Kobbekaduwa V, Tippin S, Miller G, Dinh E, Foster E, Tsao JI (2022) Northward Expansion of Amblyomma americanum (Acari: Ixodidae) into Southwestern Michigan. J Med Entomol 59:1646–1659. 10.1093/JME/TJAC082

Goddard J, Varela-Stokes AS (2009) Role of the lone star tick, Amblyomma americanum (L.), in human and animal diseases. Vet Parasitol 160:1–12. 10.1016/J.VETPAR.2008.10.089

González J, Fonseca DM, Toledo A (2023) Seasonal Dynamics of Tick Species in the Ecotone of Parks and Recreational Areas in Middlesex County (New Jersey, USA). Insects 2023, Vol 14, Page 258 14:258. 10.3390/INSECTS14030258

Guillot C, Bouchard C, Buhler K, Pelletier R, Milord F, Leighton P (2023) Quality over quantity in active tick surveillance: Sentinel surveillance outperforms risk-based surveillance for tracking tick-borne disease emergence in southern Canada. Canada Communicable Disease Report 49:50. 10.14745/ccdr.v49i23a04

Hesselbarth MHK, Sciaini M, Nowosad J, Hanss S (2025) Landscape Metrics for Categorical Map Patterns [R package landscapemetrics version 2.2.1]. CRAN: Contributed Packages. 10.32614/CRAN.PACKAGE.LANDSCAPEMETRICS

Hijmans RJ (2025a) Spatial Data Analysis [R package terra version 1.8-80]. CRAN: Contributed Packages. 10.32614/CRAN.PACKAGE.TERRA

Hijmans RJ (2025b) Geographic Data Analysis and Modeling [R package raster version 3.6-32]. CRAN: Contributed Packages. 10.32614/CRAN.PACKAGE.RASTER

Huang CI, Kay SC, Davis S, Tufts DM, Gaffett K, Tefft B, Diuk-Wasser MA (2019) High burdens of Ixodes scapularis larval ticks on white-tailed deer may limit Lyme disease risk in a low biodiversity setting. Ticks Tick Borne Dis 10:258–268. 10.1016/J.TTBDIS.2018.10.013

Hudman DA, Sargentini NJ (2018) Prevalence of Tick-Borne Pathogens in Northeast Missouri. Mo Med 115:162

Hussain A, Varga C, Allan BF, Mateus-Pinilla N, Smith RL (2025a) Spatial distribution and clustering of medically important tick species in Illinois: Implications for tick-borne disease risk. Ticks Tick Borne Dis 16:102533. 10.1016/J.TTBDIS.2025.102533

Hussain A, Varga C, Allan BF, Mateus-Pinilla N, Smith RL (2025b) Corrigendum to “Spatial distribution and clustering of medically important tick species in Illinois: Implications for tick-borne disease risk” [Ticks and Tick-borne Diseases 16 (2025) 102533]. Ticks Tick Borne Dis 16:102558. 10.1016/J.TTBDIS.2025.102558

Illinois Data Bank (2025) Illinois Data Bank. https://databank.illinois.edu/datasets/IDB-7198073. Accessed 24 Nov 2025

Illinois Department of Natural Resources (2025) Natural Division Overview. https://dnr.illinois.gov/conservation/iwap/naturaldivisionoverview.html. Accessed 24 Nov 2025

Illinois State Geological Survey (2025) Illinois County Boundaries, Polygons and Lines | clearinghouse.isgs.illinois.edu. https://clearinghouse.isgs.illinois.edu/data/reference/illinois-county-boundaries-polygons-and-lines. Accessed 24 Nov 2025

Indiana Department of Health (2025) Health: Infectious Disease Epidemiology & Prevention Division: Dermacentor variabilis. https://www.in.gov/health/idepd/zoonotic-and-vectorborne-epidemiology-entomology/vector-borne-diseases/tick-borne-diseases/dermacentor-variabilis/#Images. Accessed 25 Nov 2025

Jongejan F, Uilenberg G (2004) The global importance of ticks. Parasitology 129:S3–S14. 10.1017/S0031182004005967

Kessler WH, De Jesus C, Wisely SM, Glass GE (2022) Ensemble Models for Tick Vectors: Standard Surveys Compared with Convenience Samples. Diseases 10. 10.3390/DISEASES10020032

Lankester MW, Scandrett WB, Golsteyn-Thomas EJ, Chilton NC, Gajadhar AA (2007) Experimental transmission of bovine anaplasmosis (caused by Anaplasma marginale) by means of Dermacentor variabilis and D. andersoni (Ixodidae) collected in western Canada. Canadian Journal of Veterinary Research 71:271

Linske MA, Williams SC (2024) Evaluation of landscaping and vegetation management to suppress host-seeking Ixodes scapularis (Ixodida: Ixodidae) nymphs on residential properties in Connecticut, USA. Environ Entomol 53:268–276. 10.1093/EE/NVAE007

Louis (2012) Social, environmental, and epidemiological determinants of lone star tick associated disease emergence in St. Louis, Missouri - ProQuest. https://www.proquest.com/docview/1313776408/fulltextPDF/E4F73E33A604467EPQ/1?accountid=14553&sourcetype=Dissertations%20&%20Theses. Accessed 25 Nov 2025

Mader EM, Ganser C, Geiger A, Harrington LC, Foley J, Smith RL, Mateus-Pinilla N, Teel PD, Eisen RJ (2021) A Survey of Tick Surveillance and Control Practices in the United States. J Med Entomol 58:1503–1512. 10.1093/JME/TJAA094

Michal V, Schmidt AM, Freitas LP, Cruz OG (2025) A Bayesian hierarchical model for disease mapping that accounts for scaling and heavy-tailed latent effects. Stat Methods Med Res 34:307–321. 10.1177/09622802241293776

Minigan JN, Hager HA, Peregrine AS, Newman JA (2018) Current and potential future distribution of the American dog tick (Dermacentor variabilis, Say) in North America. Ticks Tick Borne Dis 9:354–362. 10.1016/J.TTBDIS.2017.11.012

Mori J, Brown W, Skinner D, Schlichting P, Novakofski J, Mateus-Pinilla N (2024) An Updated Framework for Modeling White-Tailed Deer (Odocoileus virginianus) Habitat Quality in Illinois, USA. Ecol Evol 14:e70487. 10.1002/ECE3.70487

Nelder MP, Russell CB, Johnson S, Li Y, Cronin K, Cawston T, Patel SN (2022) American dog ticks along their expanding range edge in Ontario, Canada. Scientific Reports 2022 12:1 12:11063-. 10.1038/s41598-022-15009-9

Paradinas I, Illian J, Smout S (2023) Understanding spatial effects in species distribution models. PLoS One 18:e0285463. 10.1371/JOURNAL.PONE.0285463

Pebesma (2025) CRAN: Package sf. In: CRAN: Contributed Packages. https://cran.r-project.org/web/packages/sf/index.html. Accessed 24 Nov 2025

Price LE, Winter JM, Cantoni JL, Cozens DW, Linske MA, Williams SC, Dill GM, Gardner AM, Elias SP, Rounsville TF, Smith RP, Palace MW, Herrick C, Prusinski MA, Casey P, Doncaster EM, Savage JDT, Wallace DI, Shi X (2024) Spatial and temporal distribution of Ixodes scapularis and tick-borne pathogens across the northeastern United States. Parasites & Vectors 2024 17:1 17:481-. 10.1186/S13071-024-06518-9

R Core Team (2025) R: The R Project for Statistical Computing. https://www.r-project.org/. Accessed 24 Nov 2025

Rochlin I, Egizi A, Lindström A (2022) The Original Scientific Description of the Lone Star Tick (Amblyomma americanum, Acari: Ixodidae) and Implications for the Species’ Past and Future Geographic Distributions. J Med Entomol 59:412–420. 10.1093/JME/TJAB215

Rochlin I, Kenney J, Little E, Molaei G (2025) Public health significance of the white-tailed deer (Odocoileus virginianus) and its role in the eco-epidemiology of tick- and mosquito-borne diseases in North America. Parasites & Vectors 2025 18:1 18:43-. 10.1186/S13071-025-06674-6

Rynkiewicz EC, Clay K (2014) Tick community composition in Midwestern US habitats in relation to sampling method and environmental conditions. Exp Appl Acarol 64:109– 119. 10.1007/S10493-014-9798-7

Sagurova I, Ludwig A, Ogden NH, Pelcat Y, Dueymes G, Gachon P (2019) Predicted northward expansion of the geographic range of the tick vector amblyomma americanum in North America under future climate conditions. Environ Health Perspect 127. 10.1289/EHP5668

Scott Dahlgren F, Paddock CD, Springer YP, Eisen RJ, Behravesh CB (2016) Expanding Range of Amblyomma americanum and Simultaneous Changes in the Epidemiology of Spotted Fever Group Rickettsiosis in the United States. Am J Trop Med Hyg 94:35. 10.4269/AJTMH.15-0580

Sharma Y, Laison EKE, Philippsen T, Ma J, Kong J, Ghaemi S, Liu J, Hu F, Nasri B (2024) Models and data used to predict the abundance and distribution of Ixodes scapularis (blacklegged tick) in North America: a scoping review. The Lancet Regional Health - Americas 32:100706. 10.1016/j.lana.2024.100706

Short SM, Pesapane R (2024) Ixodes scapularis (Blacklegged tick). Trends Parasitol 40:529– 530. 10.1016/j.pt.2024.04.002

Sirén APK, Berube J, Clarfeld LA, Sullivan CF, Simpson B, Wilson TL (2024) Accounting for missing ticks: Use (or lack thereof) of hierarchical models in tick ecology studies. Ticks Tick Borne Dis 15. 10.1016/j.ttbdis.2024.102342

Soucy JPR, Slatculescu AM, Nyiraneza C, Ogden NH, Leighton PA, Kerr JT, Kulkarni MA (2018) High-Resolution Ecological Niche Modeling of Ixodes scapularis Ticks Based on Passive Surveillance Data at the Northern Frontier of Lyme Disease Emergence in North America. https://home.liebertpub.com/vbz 18:p235–242. 10.1089/VBZ.2017.2234

Tardy O, Acheson ES, Bouchard C, Chamberland É, Fortin A, Ogden NH, Leighton PA (2023) Mechanistic movement models to predict geographic range expansions of ticks and tickborne pathogens: Case studies with Ixodes scapularis and Amblyomma americanum in eastern North America. Ticks Tick Borne Dis 14:102161. 10.1016/J.TTBDIS.2023.102161

TickEncounter (2025) American Dog Tick – TickEncounter. https://web.uri.edu/tickencounter/species/dog-tick/. Accessed 25 Nov 2025

U.S. Geological Survey (2025) Annual National Land Cover Database | U.S. Geological Survey. In: Annual National Land Cover Database (NLCD). https://www.usgs.gov/centers/eros/science/annual-national-land-cover-database. Accessed 24 Nov 2025

Wisconsin Ticks (2025) The American Dog Tick/Wood Tick (Dermacentor variabilis) – Wisconsin Ticks and Tick-borne Diseases – UW–Madison. https://wisconsin-ticks.russell.wisc.edu/wisconsin-ticks/dermacentor-variabilis/. Accessed 25 Nov 2025

