## Supplementary file for "Bayesian spatial prediction of three medically important tick species in Illinois"

### Materials and Methods

***Covariate derivation***

For land cover, NLCD classes 41, 42, and 43 were combined as forest; class 71 represented grassland; and classes 90 and 95 were combined as wetlands. County-level percentages were calculated from raster-cell coverage within each county. Forest fragmentation was quantified from class-level patch density, mean patch area, edge density, and clumpiness using eight-neighbor connectivity. Each metric was standardized before combination. The raw fragmentation score was calculated as:

$$F = z_{ED} + z_{PD} - z_{AREA} - z_{CLUMPY}$$

and then rescaled to 0-1, so that larger values represented greater fragmentation.

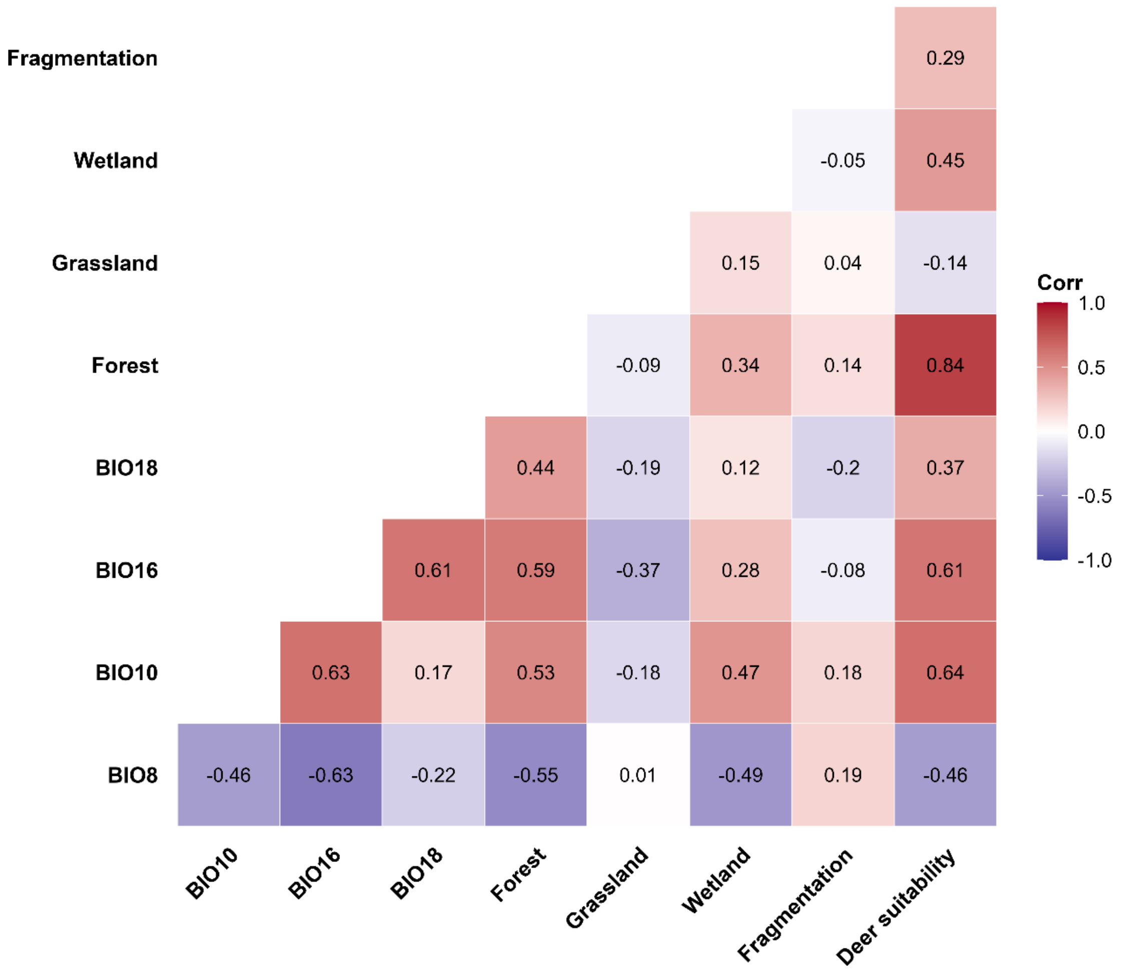


**Figure S1.** Pairwise Pearson correlations among the environmental and ecological covariates used in the analysis.

***Spatial-model implementation***
Fixed-effect coefficients were assigned *Normal (0, 1)* priors, and the intercept a Normal (0, $10^2$) prior. Penalized-complexity priors were used for the BYM2 precision [pc.prec, parameters (1, 0.01)] and mixing parameter φ [pc, parameters (0.5, 2/3)]; otherwise, default INLA settings were retained.
All retained climate principal components, land-cover principal components, and deer habitat suitability values were standardized before model fitting. The beta-binomial models were fitted using the betabinomial family in R-INLA. Model computations requested DIC, WAIC, CPO, and PIT values.

**Table S1.** Candidate model structures were fitted separately for each tick species.

| Model | Fixed effects | Spatial effect |
|---|---|---|
| M1 | Climate PC1-PC2 + land-cover PC1-PC3 + deer suitability | None |
| M2 | None | BYM2 |
| M3a | Climate PC1-PC2 | BYM2 |
| M3b | Land-cover PC1-PC3 | BYM2 |
| M3c | Deer habitat suitability | BYM2 |
| M3d | Climate PC1-PC2 + land-cover PC1-PC3 + deer suitability | BYM2 |

***Spatial cross-validation***
Balanced five-fold spatial partitions were generated independently for each species from counties with available responses. Projected county coordinates were standardized before geographic clustering. Initial folds were generated using k-means clustering with 100 random starts. Target fold sizes were defined so that folds differed by no more than one county. When initial clusters exceeded the target size, counties were iteratively reassigned from oversized to undersized folds using the reassignment that minimized the increase in squared geographic distance to the receiving cluster's centroid. Centroids were recalculated following each reassignment. Final fold sizes were 10, 10, 10, 10, and 9 counties for *A. americanum*; 16, 15, 15, 15, and 15 for *D. variabilis*; and 6 counties in each fold for *I. scapularis*. For every fold, held-out responses were set to missing, the model was refitted using the remaining observations, and posterior mean relative abundance was extracted for the withheld counties. For each held-out county, 95% posterior predictive intervals were generated from the beta-binomial posterior predictive distribution using the county's original total focal-tick count. Empirical coverage was calculated as the proportion of observed relative-abundance values falling within these intervals.

### Results

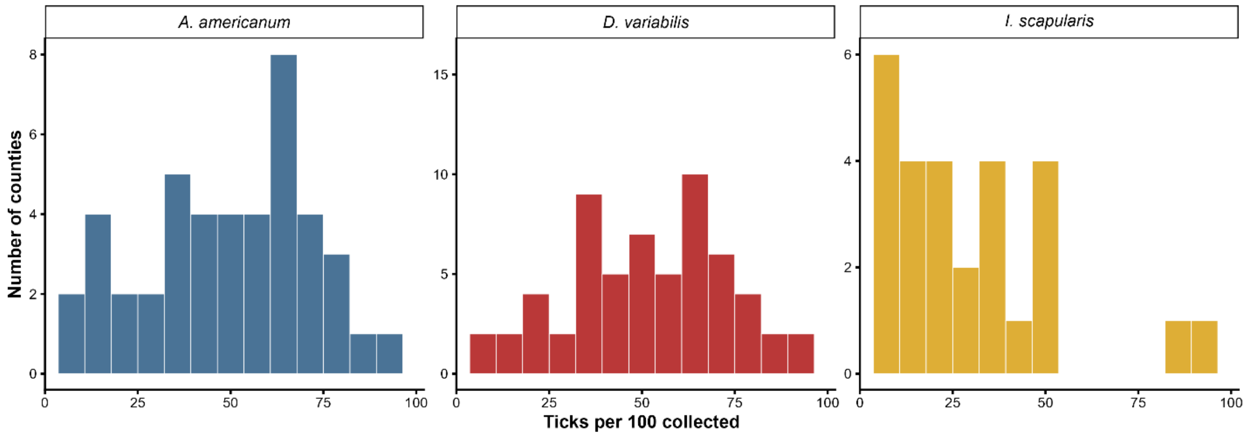


**Figure S2**. Distribution of observed county-level relative abundance for *Amblyomma americanum*, *Dermacentor variabilis*, and *Ixodes scapularis* in Illinois. Bars indicate the number of counties within each relative-abundance interval, expressed as ticks of the focal species per 100 focal ticks collected. Only counties with available species-specific observations are included.

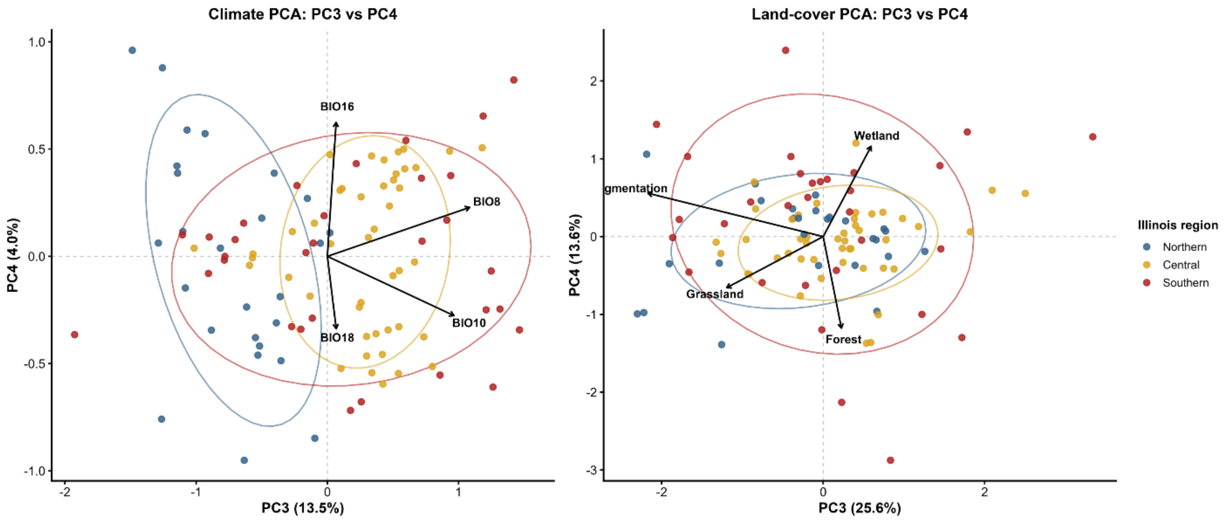


**Figure S3.** Principal component scores for PC3 and PC4 from the climate and land-cover PCAs across Illinois counties. Counties are grouped by northern, central, and southern Illinois to illustrate regional variation along the secondary environmental gradients.

**Table S2.** Principal component loadings for the climate and land-cover variables used in the PCA.

| Variable | PC1 | PC2 | PC3 | PC4 |
|---|---|---|---|---|
| ***Climate-PCA*** | | | | |
| BIO8 | -0.49 | 0.348 | 0.746 | 0.287 |
| BIO10 | 0.481 | -0.456 | 0.663 | -0.348 |
| BIO16 | 0.608 | 0.11 | 0.046 | 0.785 |
| BIO18 | 0.398 | 0.812 | 0.046 | -0.424 |
| ***Land-cover PCA*** | | | | |
| Forest cover | -0.693 | 0.336 | 0.088 | -0.632 |
| Grassland cover | -0.126 | -0.801 | -0.467 | -0.353 |
| Wetland cover | -0.684 | -0.302 | 0.232 | 0.622 |
| Forest fragmentation | -0.19 | 0.393 | -0.848 | 0.299 |

**Note:** *BIO8, mean temperature of the wettest quarter; BIO10, mean temperature of the warmest quarter; BIO16, precipitation of the wettest quarter; BIO18, precipitation of the warmest quarter. Loadings are shown to three decimal places.*

**Table S3.** Model-comparison statistics for candidate beta-binomial models of county-level relative abundance of three tick species in Illinois. Lower WAIC, DIC, and LCPO values indicate better model support. The model selected for each species is shown in bold.

| Tick species | Model | WAIC | DIC | LCPO |
|---|---|---|---|---|
| *A. americanum* | M1: Environmental-only | 252.712 | 252.776 | 126.258 |
| *A. americanum* | M2: BYM2 spatial-only | 254.899 | 255.468 | 125.379 |
| *A. americanum* | M3a: Climate PC1–PC2 + BYM2 | 259.429 | 259.098 | 129.328 |
| *A. americanum* | M3b: Land-cover PC1–PC3 + BYM2 | 262.168 | 260.651 | 129.864 |
| *A. americanum* | M3c: Deer habitat suitability + BYM2 | 253.536 | 253.612 | 126.617 |
| ***A. americanum*** | **M3d: Climate + land cover + deer + BYM2** | **250.922** | **251.124** | **124.816** |
| *D. variabilis* | M1: Environmental-only | 330.299 | 330.435 | 164.276 |
| ***D. variabilis*** | **M2: BYM2 spatial-only** | **320.137** | **320.411** | **156.634** |
| *D. variabilis* | M3a: Climate PC1–PC2 + BYM2 | 330.998 | 330.387 | 165.31 |
| *D. variabilis* | M3b: Land-cover PC1–PC3 + BYM2 | 327.095 | 328.235 | 162.46 |
| *D. variabilis* | M3c: Deer habitat suitability + BYM2 | 327.557 | 328.423 | 163.309 |
| *D. variabilis* | M3d: Climate + land cover + deer + BYM2 | 328.724 | 328.335 | 162.678 |
| *I. scapularis* | M1: Environmental-only | 140.513 | 140.006 | 102.857 |
| *I. scapularis* | M2: BYM2 spatial-only | 139.303 | 139.502 | 71.634 |
| ***I. scapularis*** | **M3a: Climate PC1–PC2 + BYM2** | **137.935** | **137.482** | **69.087** |
| *I. scapularis* | M3b: Land-cover PC1–PC3 + BYM2 | 139.899 | 139.727 | 70.376 |
| *I. scapularis* | M3c: Deer habitat suitability + BYM2 | 139.847 | 139.191 | 69.773 |
| *I. scapularis* | M3d: Climate + land cover + deer + BYM2 | 138.734 | 137.683 | 143.695 |

***Note:*** *WAIC, Watanabe-Akaike information criterion; DIC, deviance information criterion; LCPO, leave-one-out conditional predictive ordinate summary.*

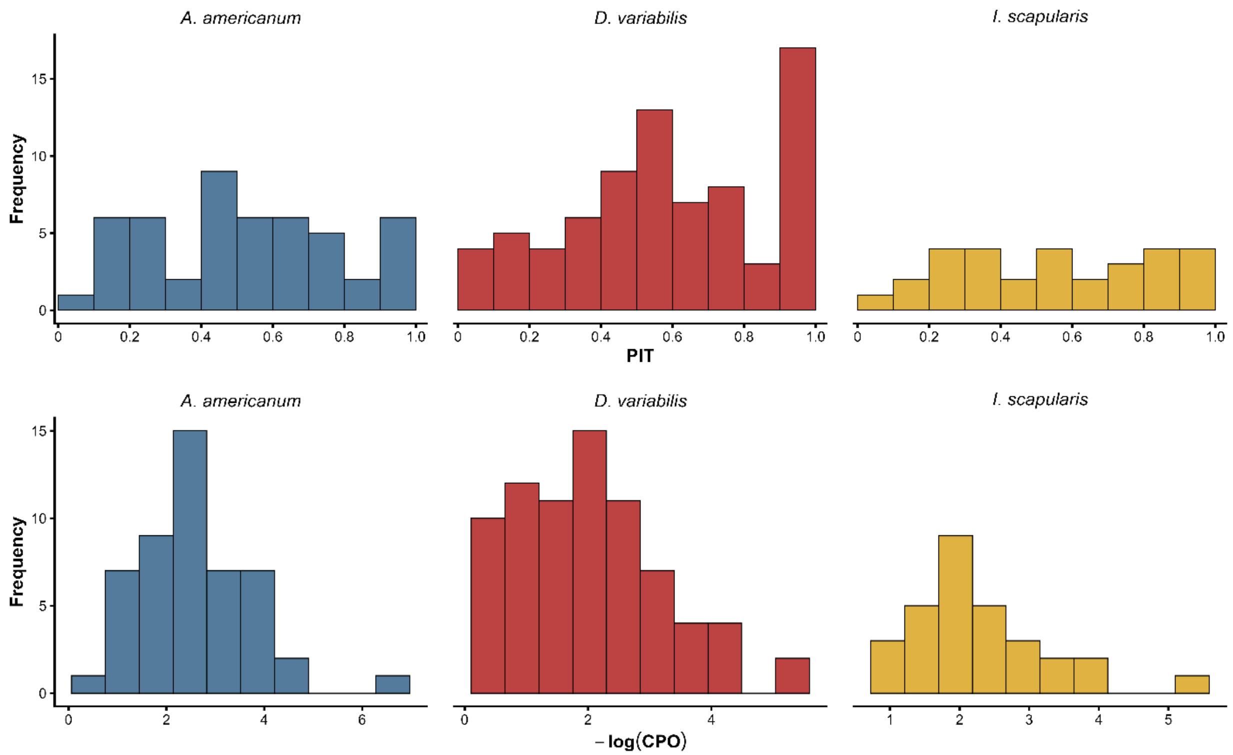


**Figure S4.** Predictive diagnostics for the selected species-specific Bayesian models. The upper panels show probability integral transform (PIT) distributions, and the lower panels show distributions of −log(CPO) for *A. americanum*, *D. variabilis*, and *I. scapularis*.

**Table S4.** Spatial cross-validation performance of spatial-only and environmental-plus-spatial models for the three tick species.

| Species | Model Type | MSE | RMSE | MAE | Bias |
|---|---|---|---|---|---|
| *A. americanum* | Spatial-only | 0.085893 | 0.293075 | 0.237559 | -0.04491 |
| *A. americanum* | Environmental + spatial | 0.07682 | 0.277165 | 0.211607 | -0.00864 |
| *D. variabilis* | Spatial-only | 0.066979 | 0.258803 | 0.21959 | -0.02106 |
| *D. variabilis* | Environmental + spatial | 0.078629 | 0.280408 | 0.223378 | -0.09641 |
| *I. scapularis* | Spatial-only | 0.065132 | 0.255209 | 0.198609 | 0.015633 |
| *I. scapularis* | Environmental + spatial | 0.055931 | 0.236497 | 0.183772 | 0.011341 |

***Note:*** *Performance was evaluated using balanced five-fold spatial cross-validation. MSE, mean squared error; RMSE, root mean squared error; MAE, mean absolute error; Bias, mean prediction bias. Lower MSE, RMSE, and MAE indicate better predictive performance, while bias values closer to zero indicate less systematic prediction error.*

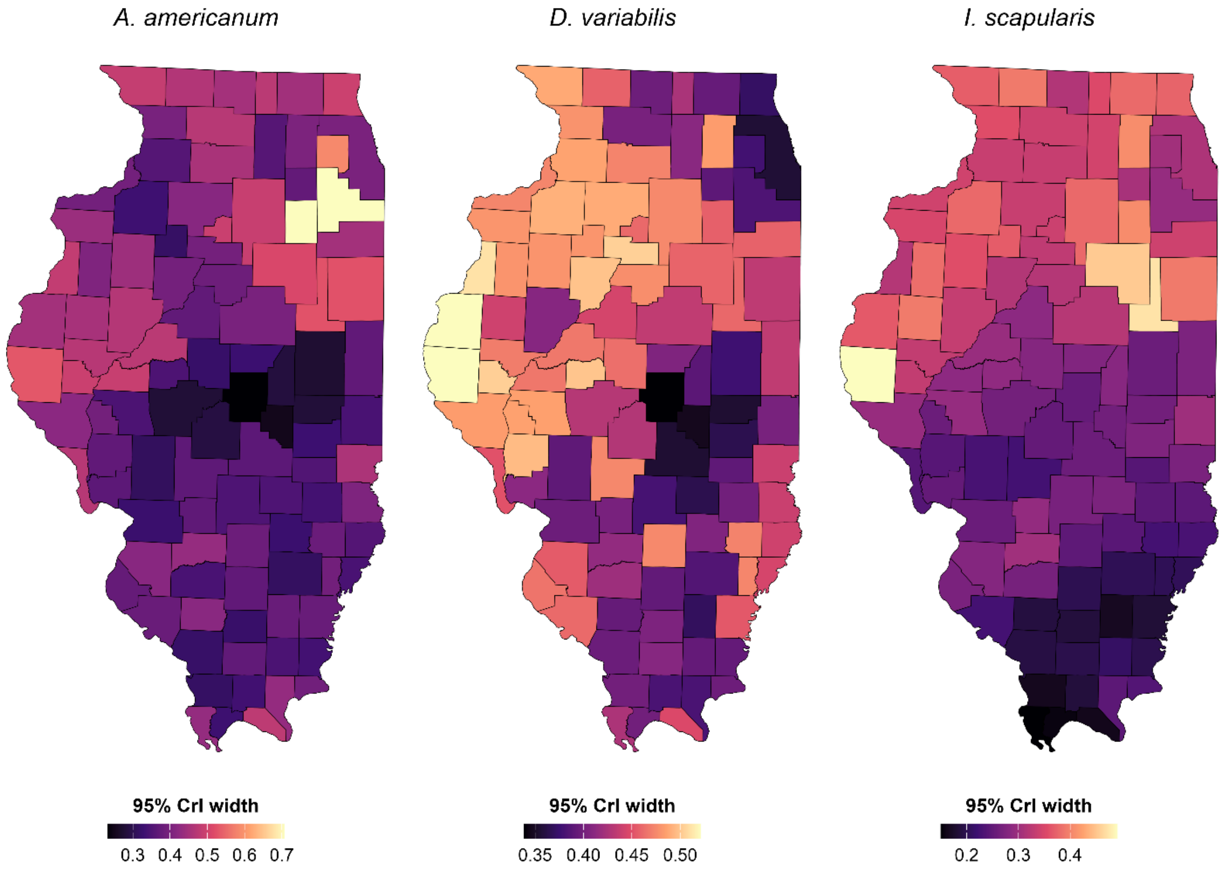


**Figure S5.** Spatial distribution of posterior uncertainty in predicted relative abundance for *Amblyomma americanum*, *Dermacentor variabilis*, and *Ixodes scapularis*, expressed as the width of the 95% credible interval. Wider intervals indicate greater posterior uncertainty.
